# Potential Application of Bacteriophages in Enrichment Culture for Improved Prenatal *Streptococcus agalactiae* Screening

**DOI:** 10.1101/384222

**Authors:** Jumpei Uchiyama, Hidehito Matsui, Hironobu Murakami, Shini-chiro Kato, Naoki Watanabe, Tadahiro Nasukawa, Keijiro Mizukami, Masaya Ogata, Masahiro Sakaguchi, Shigenobu Matsuzaki, Hideaki Hanaki

**Affiliations:** School of Veterinary Medicine, Azabu University, Kanagawa, Japan; Kitasato Institute for Life Sciences, Kitasato University, Tokyo, Japan; Research Institute of Molecular Genetics, Kochi University, Kochi, Japan; Kochi Medical School, Kochi University, Kochi, Japan

**Keywords:** phage, *Enterococcus faecalis*, *Streptococcus agalactiae*, culture enrichment

## Abstract

Vertical transmission of *Streptococcus agalactiae* can cause neonatal infections. A culture test in the late stage of pregnancy is used to screen for the presence of maternal *S. agalactiae* for intrapartum antibiotic prophylaxis. For the test, vaginal-rectal swab sampling is immediately followed by enrichment culture and bacterial identification. In some cases, *Enterococcus faecalis* competes with and overgrowths *S. agalactiae* in the enrichment culture. Consequently, the identification test occasionally yields false-negative results. Bacterial viruses, bacteriophages (phages), infect and kill specific host bacteria. In the current study, we explored the feasibility of using phages to minimize the undesirable *E. faecalis* outgrowth and facilitate *S. agalactiae* detection in an experimental setting. Phage mixture was prepared using three phages that specifically infect *E. faecalis:* phiEF24C, phiEF17H, and phiM1EF22. The mixture inhibited the growth of 86.7% (26/30) of *E. faecalis* strains tested in the enrichment broth. When single strains of *E. faecalis* and *S. agalactiae* were inoculated in the enrichment broth containing the phage mixture, bacterial growth was inhibited or facilitated, respectively. Further, several sets of *S. agalactiae* and *E. faecalis* strains were co-cultured, and bacteria were detected on chromogenic agar after the enrichment culture. *S. agalactiae* was dominant after plating a phage mixture-treated co-culture, while it was barely detected after plating the untreated co-culture. Considering these observations, the phage mixture can be employed in the *S. agalactiae* culture test to increase test accuracy.

## INTRODUCTION

*Streptococcus agalactiae* (also called group B streptococcus) is vertically transmitted to the newborn during delivery, and can cause neonatal infections (1, 2). Common early-onset diseases caused by this organism in infants include sepsis and pneumonia, and (rarely) meningitis (1, 2). To prevent such infections, prenatal *S. agalactiae* culture test is recommended in the late stage of pregnancy (1, 2). In the case of a positive test result, the pregnant carrier is prophylactically treated with antibiotics to prevent vertical transmission of *S. agalactiae* during the intrapartum period (1, 2).

For the *S. agalactiae* culture test, the Center of Disease Control and Prevention highly recommends an enrichment culture, followed by conventional *S. agalactiae* identification (3, 4). In the culture test, a swab is taken from the vaginal and anorectal areas, and the samples are inoculated and cultured in an enrichment culture broth selective for *S. agalactiae*. After the enrichment culture, bacterial identification is performed, e.g., by using the Christie-Atkins-Munch-Petersen test, serologic identification, growth on chromogenic agar, and nucleic acid amplification (4). However, although a selective culture broth is used for the enrichment culture, *S. agalactiae* is poorly recovered along with overgrowth of *Enterococcus faecalis* (5-8). This may lead to false-negative results in the subsequent identification tests (5-8). To address the problem of false-negative results, selective antimicrobial agents to be included in the enrichment broth should be re-evaluated.

Bacteriophages (phages), i.e., bacterial viruses, infect specific bacteria. Some phages infect and lyse bacteria at the specificity level of species and strains. These phage characteristics have been used to eliminate most cells in a bacterial population and facilitate the isolation of less prevalent environmental bacteria that produce novel bioactive compounds (9). Phage applicability for the isolation of food-poisoning microbes in the food microbiology field has also been examined (10). Hence, potentially, phage application might also be used to reduce the unwanted growth of *E. faecalis* and facilitate *S. agalactiae* growth in an *S. agalactiae* enrichment culture in clinical microbiology. Indeed, phages that specifically infect *E. faecalis* have been isolated from environmental samples, such as sewage and canal water (11-13). In the current study, we examined the applicability of *E. faecalis-specific* phages to suppress *E. faecalis* growth in *S. agalactiae* enrichment culture.

## MATERIALS AND METHODS

### Bacteria, phages, and culture media

Strains of *E. faecalis* (*n* = 30), *S. agalactiae* (*n* = 7), *Enterococcus avium* (*n* = 5), and *Enterococcus faecium* (*n* = 5) were isolated from vaginal swabs using the Chrom-ID Strepto B test (bioMérieux, Marcy-l’Étoile, France). The swabs were obtained after random sampling at local hospitals in eastern Japan (Table S1). Bacteria were cultured at 37°C under aerobic or microaerobic (i.e., 5% CO_2_) condition, as appropriate, based on their specific growth requirements (Table S1).

Phage phiEF24C has been isolated and characterized, as described elsewhere (12,14,15). Phage phiEF17H was newly isolated from canal water in Kochi (Japan). Phage phiM1EF22 was newly isolated from sewage water in Tokyo (Japan) (Table S2). The isolation procedures are described elsewhere (12). *E. faecalis* strains KUEF01, KUEF25, and KUEF27, described in Table S1, were used as host bacteria for phages phiEF24C, phiEF17H, and phiM1EF22, respectively, for phage amplification and plaque assay. Bacterial-phage suspensions were cultured aerobically at 37°C.

*Enterococcus* spp. and phages were cultured in tryptic soy broth or agar (TSA), and *S. agalactiae* was cultured in Todd-Hewitt broth (THB), unless stated otherwise. Granada-type broth with slight modification [GBwSM; 25.0 g/l proteose peptone no. 3, 14.0 g/l soluble starch, 2.5 g/l glucose, 1.0 g/l pyruvic acid sodium salt, 0.1 g/l cysteine hydrochloride, 0.3 g/l magnesium sulfate, 11.0 g/l 3-(*N*-morpholino)propane sulfonic acid, 10.7 g/l disodium hydrogen phosphate, 0.5 mg/l crystal violet, 10 mg/l colistin sulfate, 10 mg/l metronidazole, and 15 mg/l nalidixic acid, pH 7.4] was originally prepared as the *S. agalactiae* enrichment broth (16, 17). Alternatively, the pigmented enrichment Lim broth (modified Lim broth; Kyokuto Pharmaceutical Industrial, Tokyo, Japan) was used as an *S. agalactiae* enrichment broth. Unless stated otherwise, all culture media were purchased from Becton, Dickinson, and Co. (Franklin Lakes, NJ). All chemicals and reagents were purchased from Nacalai Tesque (Kyoto, Japan) and FUJIFILM Wako Pure Chemical (Osaka, Japan).

### Phage genome sequencing

After phage amplification, phage particles were purified from 500 ml of phage lysate by CsCl density-gradient centrifugation, as described elsewhere (18). Phage genomic DNA was then prepared by phenol-chloroform extraction of the collected purified phage band, as described (18). Shotgun library was prepared for each phage DNA using the GS FLX Titanium rapid library preparation kit (Roche Diagnostics, Indianapolis, IN), according to the manufacturer’s instruction. The libraries were analyzed using a GS Junior 454 sequencer (Roche Diagnostics). The sequence reads were assembled using the 454 Newbler software (version 3.0; 454 Life Sciences, Branford, CT). The genome sequences were analyzed by using BLASTn at the National Center for Biotechnology Information (NCBI; https://blast.ncbi.nlm.nih.gov/Blast.cgi?PROGRAM=blastn&PAGE_TYPE=BlastSearch&LINK_LOC=blasthome; last accessed: 5 May, 2018). The genomes were annotated using a prokaryotic genome annotation pipeline, DFAST (https://dfast.nig.ac.jp/) (19, 20).

### Multi-locus sequence typing (MLST) of *E. faecalis* strains

*E. faecalis* strains were cultured overnight, bacterial DNA was extracted, and MLST analysis was performed, according to the procedures described elsewhere (21). The sequence alleles were analyzed using the *E. faecalis* MLST database (https://pubmlst.org/efaecalis/; last accessed: 5 January, 2018) to designate sequence types (STs) (22). The concatenating allele sequences were analyzed using MEGA 7.0.18, and sequence alignment implemented in ClustalW was followed by phylogenetic tree construction by the UPGMA method (23).

### Examination of antibacterial activity of *E. faecalis* to *S.agalactiae*

The anti-*S*. *agalactiae* activity of *E. faecalis* was examined by spot-on-lawn assay, as described elsewhere (24). Briefly, 200 μl of overnight bacterial culture of a single *S. agalactiae* strain were mixed with a melted 0.5% (w/v) soft agar and plated onto 1.5% (w/v) agar. One microliter of *E. faecalis* overnight culture was spotted on the solidified top agar. After incubation overnight at 37°C in a microaerophilic condition, *S. agalactiae* growth around the spotted *E. faecalis* was examined.

### Analysis of phage lytic activity

The phage host range was determined by a streak test, as described elsewhere (12, 15). Briefly, 200 μl of overnight bacterial culture of a single bacterial strain were mixed with a melted 0.5% (w/v) soft agar and plated onto 1.5% (w/v) agar. Phage suspension (ca. 1.0 × 10^8–9^ PFU/ml) was streaked onto the solidified top agar. After incubation overnight at 37°C, bacterial lysis, with or without plaque formation, was examined.

### Analysis of bacterial growth inhibition by the phage mixture in *S. agalactiae* enrichment broths

Bacteria were cultured until optical density of 0.4–0.6 at 600 nm. After washing and diluting with GBwSM, 3.0–5.0 × 10^8^ CFU/ml bacterial suspension was prepared in GBwSM. Each phage suspension was diluted in THB to ca. 3.0 × 10^8^ PFU/ml, 3.0 × 10^6^ PFU/ml, and 3.0 × 10^4^ PFU/ml. Phage mixtures were prepared by mixing equal volumes of phage suspensions at the same dilution. Then, 5 μl of bacterial suspension and 5 μl of phage mixture were added to 140 μl of GBwSM in a well of a flat-bottomed polystyrene 96-well plate (AS ONE Co., Osaka, Japan). In the experiments, 5 μl of THB was used instead of the bacterial suspension and/or phage mixture as a control. The 96-well plate was incubated at 37°C and sample turbidity was measured over time at 595 nm, using a Multiskan JX spectrophotometer (Thermo Labsystems, Stockholm, Sweden). The experiments were conducted in triplicate, and the growth curves were then plotted using averaged values with standard deviations.

### Analysis of bacterial densities in *S.agalactiae* and *E. faecalis* co-culture with phage mixtures

Rifampicin-resistant mutant clone of *S. agalactiae* was isolated by aerobically culturing *S. agalactiae* strain KUGBS2 on TSA containing 20 μg/ml rifampicin at 37°C for 2 d. The putative mutant clones were re-purified at least three times; each re-purification round was repeated for 1 d under the same incubation conditions. One resultant rifampicin-resistant mutant clone of strain KUGBS2 was obtained and was tentatively designated as strain KUGBS2rif. *S. agalactiae* strain KUGBS2rif and *E. faecalis* strain KUEF08 were cultured individually until optical density of 0.4–0.6 at 600 nm. After diluting with the enrichment broth, suspensions of 3.0 × 10^4^ CFU/ml *S. agalactiae* strain KUGBS2rif and 3.0 × 10^7^ CFU/ml *E. faecalis* strain KUEF08 were prepared. Each phage suspension was diluted with THB to ca. 3.0 × 10^6^ PFU/ml or 3.0 × 10^4^ PFU/ml. By mixing equal volumes of phage suspensions at the same dilution, mixtures of two different dilutions of phages were prepared.

For the experiment, 100 μl each of *S. agalactiae* strain KUGBS2rif and *E. faecalis* strain KUEF08, and 300 μl of phage mixture were added to 10 ml of the *S. agalactiae* enrichment broth. As negative controls, the same volume of THB was added instead of bacterial suspensions and/or phage suspensions. The mixtures were microaerobically incubated at 37°C for 24 h. Total bacterial density, and *S. agalactiae* strain KUGBS2rif and *E. faecalis* strain KUEF08 densities were determined. Total bacterial densities were determined on TSA. TSA supplemented with 20 μg/ml rifampicin and *Enterococcus-selective* agar (EF agar base “Nissui”; Nissui Pharmaceutical Co., Tokyo, Japan) were used to determine the densities of *S. agalactiae* strain KUGBS2rif and *E. faecalis* strain KUEF08, respectively. *S. agalactiae* strain KUGBS2rif did not grow on the *Enterococcus-selective* agar; conversely, *E. faecalis* strain KUEF08 did not grow on TSA containing 20 μg/ml rifampicin.

### Detection of bacteria on chromogenic selective agar after *S. agalactiae* and *E. faecalis* co-culture with phage mixtures

*S. agalactiae* and *E. faecalis* were cultured individually until optical density of 0.4–0.6 at 600 nm. *S. agalactiae* and *E. faecalis* cultures were diluted with THB to ca. 3.0–5.0 × 10^4^ CFU/ml and ca. 3.0 × 10^7^ CFU/ml, respectively. After dilution of individual phage suspensions in THB to ca. 1.0 × 10^7^ PFU/ml, phage mixture was prepared by mixing equal volumes of the diluted phage suspensions.

For the experiment, 30 μl each of bacterial suspensions of *S. agalactiae* and *E. faecalis*, and 30 μl of phage mixture were added to 3 ml of the enrichment broth. As a negative control, the same volume of THB was added instead of the phage mixture. After 24-h incubation at 37°C, a loop-full of the suspension was inoculated on the Chrom-ID Strepto B agar (bioMérieux). After 24-h incubation at 37°C in darkness, colony color and appearance on agar plates were examined. All incubations were made under microaerophilic conditions.

### Accession numbers

The phiEF17H and phiM1EF22 genome sequences were deposited in the GenBank under the accession numbers AP018714 and AP018715, respectively.

### Statistical analysis

The data were statistically analyzed using EZR (Saitama Medical Center, Jichi Medical University, Saitama, Japan), which is a graphical user interface for R (The R Foundation for Statistical Computing, Vienna, Austria) (25). Student’s *t*-tests were used to analyze differences between bacterial densities in different treatments. The value of P < 0.01 was considered to indicate statistically significant difference.

### Data availability

By publishing in the journal, the authors agree that, subject to requirements or limitations imposed by local and/or U.S. Government laws and regulations, any materials and data that are reasonably requested by others are available from a publicly accessible collection or will be made available in a timely fashion, at reasonable cost, and in limited quantities to members of the scientific community for noncommercial purposes. The authors guarantee that they have the authority to comply with this policy either directly or by means of material transfer agreements through the owner.

## RESULTS AND DISCUSSION

### Phage characteristics

Phages phiEF24C, phiEF17H, and phiM1EF22 were used in the current study. Phage phiEF24C, one of the well-studied *Enterococcus* phages, is classified into the family *Myoviridae* subfamily *Spounavirinae* (26). Based on the whole-genome sequence similarities to the phage phiEF24C genome, phages phiEF17H and phiM1EF22 share viral taxonomy with that phage (Table S3). Phages sharing this particular viral taxonomy are highly virulent toward host bacteria (27). MLST analysis of the *E. faecalis* vaginal swab isolates (Fig. 1) revealed that 43.3% (13/30) of *E. faecalis* strains were phylogenetically closely related, representing either ST16 or ST179. The remaining strains [56.7% (17/30)] were genetically diverse.

**FIG 1.**
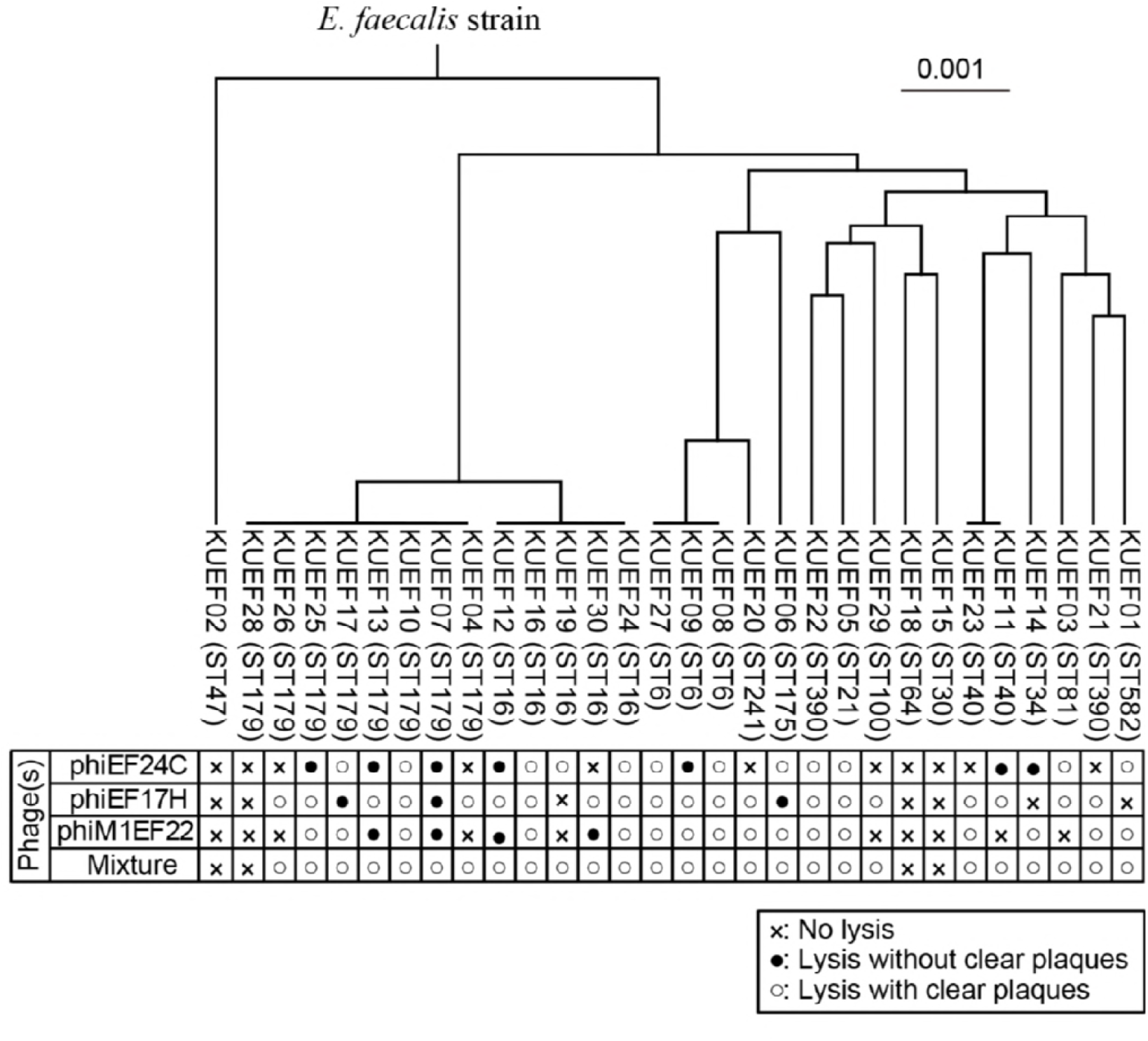
*E. faecalis* strains isolated from vaginal swabs and their sensitivity to phages. Phylogenetic tree of *E. faecalis* strains was constructed based on the concatenated MLST alleles. In the phylogenetic tree, *E. faecalis* strain names are followed by STs in brackets. Phage sensitivities to each phage and phage mixture are shown below the phylogenetic tree.

Moreover, *E. faecalis* strains, which interfere *S. agalactiae* culture tests, do not seem to show antibacterial activity (e.g., bacteriocin production) to the *S. agalactiae*, while some of them are able to show anti-bacterial activity to a variety of bacteria (5, 28). Testing the antibacterial activity to *S. agalactiae* strains by the spot-on-lawn assay, no anti-*S*. *agalactiae* activity was observed among these *E. faecalis* strains.

Phage lytic activity was examined by a streak test using these *E. faecalis* strains (Fig. 1). Phages phiEF24C, phiEF17H, and phiM1EF22 showed lytic activity toward 63.3% (19/30), 76.7% (23/30), and 66.7% (20/30), respectively, of the tested *E. faecalis* strains. Lytic activities of the three phages with other bacterial vaginal swab isolates (*E. avium, E. faecium*, and *S. agalactiae* strains) were also examined, but no lytic activity was observed.

### Lytic activity of the phage mixture as an *E. faecalis*-selective antimicrobial agent

Phages phiEF24C, phiEF17H, and phiM1EF22 lysed different *E. faecalis* strains and also some common strains. Theoretically, a combination of these three phages lysed a broader range of *E. faecalis* strains than any single phage tested. Phage mixture containing the three phages was prepared by mixing phage particles in 1:1:1 ratio. The lytic spectrum of the phage mixture was then examined by using a streak test. The phage mixture showed lytic activity toward 86.7% (26/30) of *E. faecalis* strains tested (Fig. 1). Four *E. faecalis* strains were not lysed by the phages, namely, KUEF02 (MLST ST47), KUEF18 (ST64), KUEF15 (ST30), and KUEF28 (ST179). The phage mixture did not show any lytic activity with the other tested bacteria, i.e., *E. avium, E. faecium*, and *S. agalactiae*.

To evaluate the effect of the phage mixture on the growth of *E. faecalis* in the enrichment broth, *E. faecalis* growth was monitored in the presence of three dilutions of the phage mixture in GBwSM. The medium lacked the customary pigment enhancer methotrexate to improve data reliability (16, 17). After bacterial inoculation, phage mixture was added to the multiplicity of infection (MOI) of each phage of 1, 10^-2^, or 10^-4^. For the *S. agalactiae* enrichment culture, 18–24-h incubation is recommended (3, 4). Consequently, the turbidity of cultures of 30 *E. faecalis* strains described in Fig. 1 was recorded in the presence or absence of the phage mixture over 24 h.

To simplify data interpretation, *E. faecalis* growth curves were tentatively categorized into four patterns (A–D), after comparing with the growth curve of untreated *E. faecalis* (Fig. 2A). For strains representing pattern A growth curve, *E. faecalis* growth was inhibited throughout the experiment. For strains representing pattern B growth curve, *E. faecalis* growth was initially inhibited (8 h post inoculation) and then gradually recovered. For strains representing pattern C growth curve, bacteria showed an initial growth but were then lysed. Pattern C was a typical phage lysis pattern at the lower MOI tested. *E. faecalis* strains that exhibited growth patterns A–C were inhibited by the phage treatment. The tested *E. faecalis* strains treated with higher concentrations of phage mixtures and ones that were most sensitive to phage mixture treatments tended to exhibit growth patterns A, B, and C, in the order of strongest to weakest inhibition. On the other hand, phage treatment did not inhibit *E. faecalis* strains exhibiting growth pattern D. The four phage-insensitive *E. faecalis* strains (KUEF02, KUEF18, KUEF15, and KUEF28), and *E. avium, E. faecium*, and *S. agalactiae* strains exhibited growth pattern D. The reasons for such tentative growth pattern categorization (i.e., growth patterns A-D) were probably due to (1) different sensitivity of *E. faecalis* strains to phages, and (2) the use of three dilutions of the phage mixture in experiments.

**FIG 2.**
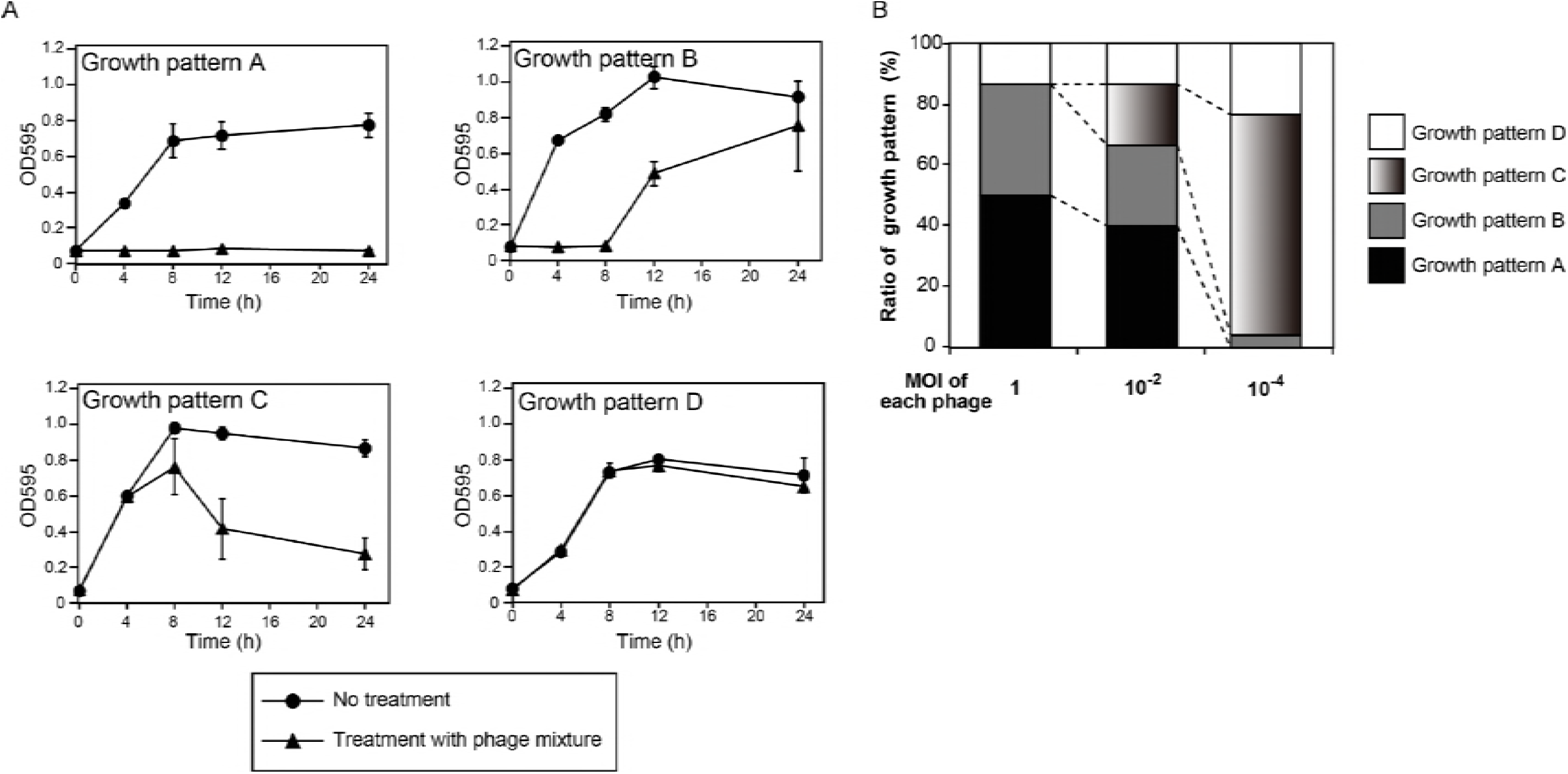
Effects of phage mixture treatment on *E. faecalis* growth in GBwSM. (A) Four growth patterns of *E. faecalis* strains treated with phage mixture. *E. faecalis* strains were treated, or not, with the phage mixture, and culture turbidities were evaluated by measuring optical density at 595 nm (OD595) over time. Thirty *E. faecalis* strains isolated from vaginal swabs, described in Fig. 1, were individually tested. The growth curves were graphed using means and standard deviations from triplicate experiments. The growth curves of phage-treated *E. faecalis* strains were categorized into four types, patterns A–D, after comparing with the growth curves of untreated *E. faecalis* strains. For the growth patterns A–C, phage treatments with all tested phage dilutions inhibited *E. faecalis* growth compared with the negative control (i.e., no phage treatment). In growth pattern D, phage mixture did not inhibit bacterial growth. The representative growth curves were presented: pattern A, strain EF08 treated with phage mixture at MOI of each phage to *E. faecalis* at 1; pattern B, strain EF22 treated with phage mixture at MOI of each phage to *E. faecalis* at 10^-2^; pattern C, strain EF05 treated with phage mixture at MOI of each phage to *E. faecalis* at 10^-4^; pattern D, strain EF02 treated with phage mixture at MOI of each phage to *E. faecalis* at 1. (B) Summary of the effect of phage mixture treatments on *E. faecalis* growth when three different phage mixture dilutions were tested. Phage densities (MOI) for each phage type were 1, 10^-2^, or 10^-4^. *E. faecalis* growth patterns were classified based on the *E. faecalis* growth patterns described in Fig. 2A, and are summarized as cumulative bar graphs.

The tentative grouping of growth patterns of the 30 tested *E. faecalis* is summarized in Fig. 2B. Phage mixture treatments at the highest and the second highest phage density (i.e., MOI 1 and 10^-2^, respectively) inhibited the growth of 86.7% (26/30) of *E. faecalis* strains, which was in agreement with the results of the streak test (Fig. 1). On the other hand, when the phage density was reduced (i.e., MOI 10^-4^), the number of *E. faecalis* strains whose growth was inhibited was reduced (76.7%, 23/30 strains). The latter 23 strains largely represented growth pattern C, probably because the input phage titer was much lower than that in the other phage mixture treatments. The seven remaining *E. faecalis* strains that exhibited growth pattern D included the four phage mixture-insensitive strains mentioned above and strains KUEF03 (ST3), KUEF19 (ST16), and KUEF30 (ST16). In the current study, a correlation between growth patterns and STs was not observed among the *E. faecalis* strains tested.

Based on these observations, we concluded that the phage mixture inhibited the growth of 86.7% (26/30) of *E. faecalis* strains tested, when used at sufficiently high density (i.e., MOI of at least 10^-2^).

### The effect of phage mixture on *S. agalactiae* and *E. faecalis* cell densities in experimental enrichment cultures

Phage mixture may have been contaminated with this anti-*S*. *agalactiae* agents during phage mixture preparation (i.e., during phage propagation on the *E. faecalis* host). However, incubation of phage mixture with *S. agalactiae* did not significantly affect bacterial viability, compared with a THB-treated negative control (Fig. S1), excluding the possibility of phage mixture contamination with anti-*S*. *agalactiae* substances.

The effects of phage mixture were then examined in co-cultures of *E. faecalis* and *S. agalactiae* in the Granada-type enrichment broth. The rifampicin-resistant mutant clone of strain KUGBS2, KUGBS2rif, was isolated. *S. agalactiae* strain KUGBS2rif and *E. faecalis* strain KUEF08 were used to quantify viable bacterial individually. To mimic the situation that *S. agalctaie* was poorly recovered, *S. agalactiae* strain KUGBS2rif and *E. faecalis* strain KUEF08 were inoculated into GBwSM at 3.0 × 10^2^ CFU/ml and 3.0 × 10^5^ CFU/ml, respectively. Either of the two dilutions of phage mixture (at MOI of each phage of 10^-1^ and 10^-3^ to *E. faecalis*) were added. As a negative control, THB was used instead of the phage mixture.

Changes in bacterial cell density (total bacteria, *S. agalactiae*, and *E. faecalis*) were then monitored over time (Fig. 3). Based on the determined total bacteria numbers, bacteria grew exponentially for up to 12 h, following which the cultures entered the stationary phase of growth. Hence, 12–24-h incubation was sufficient to achieve bacterial enrichment in that particular experimental setting. Moreover, changes in *E. faecalis* and *S. agalactiae* cell densities were then evaluated. In the negative control group (i.e., no phage treatment), *E. faecalis* grew much better than *S. agalactiae*. In the phage treatment groups, the opposite was observed. After 12 and 24 h, in the phage treatment groups, cell density of *E. faecalis* was significantly lower than that of *S. agalactiae* (P < 0.01).

**FIG 3.**
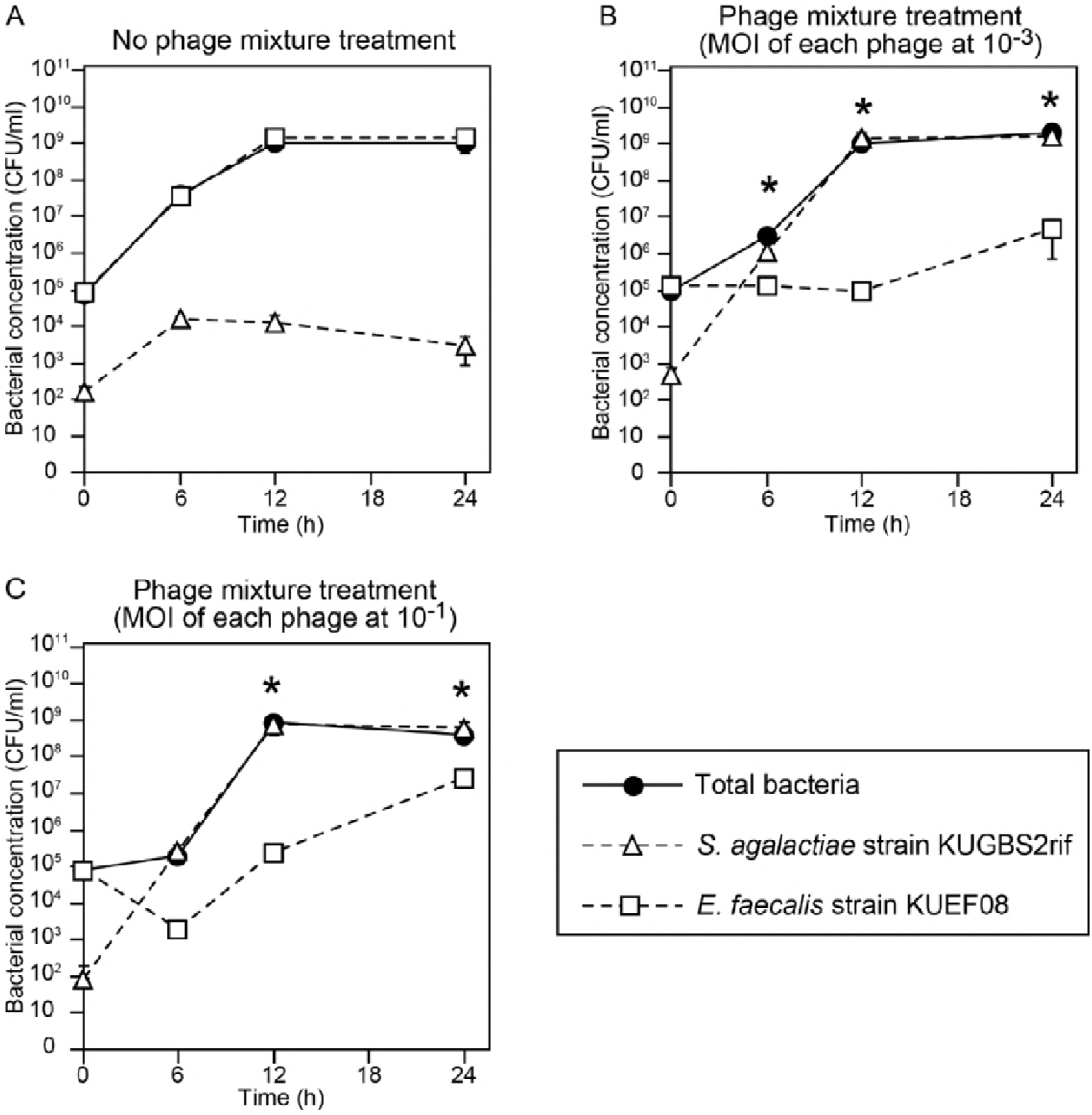
Growth of *E. faecalis* and *S. agalactiae* co-cultures in the presence or absence of phage mixtures in GBwSM. No phage treatment (A); or treatment with phages at 10^-3^ at (B) or 10^-1^ (C) MOI of each phage to *E. faecalis*. The means with standard deviations were calculated from triplicate experiments, and are plotted as points with error bars. Time points at which *S. agalactiae* density was significantly higher than that of *E. faecalis* are indicated by asterisks (P < 0.01; Student’s *t*-test).

Several types of enrichment broths are commercially available for *S. agalactiae* culture test. In addition to the GBwSM medium, we also evaluated the effectiveness of phage treatment of an *E. faecalis* and *S. agalactiae* co-culture in the commercially available pigmented enrichment Lim broth (Fig. S2). The experiments were performed as described above. Inhibition of *E. faecalis* growth, compared with the untreated group, was observed in phage treatment groups at both MOIs of each phage tested (i.e., 1 and 10^-2^) (Fig. S3). This indicated that phage mixture inhibited *E. faecalis* growth and facilitated *S. agalactiae* growth in both *S. agalactiae* enrichment broths.

### Efficient detection of *S. agalactiae* after enrichment culturing in the presence of phage mixture

In the *S. agalactiae* culture test, bacteria are generally identified in a culture aliquot after enrichment culture. Consequently, we then evaluated the efficiency of *S. agalactiae* identification after the experimental enrichment culture. As the identification assay, we used growth on the *S. agalactiae* chromogenic agar, in which *S. agalactiae* colonies are distinguished from *E. faecalis* colonies based on color. We tested several *S. agalactiae–E. faecalis* combinations. *E. faecalis* and *S. agalactiae* strains were first inoculated at 100:1 ratio in the Granada-type enrichment broth, the phage mixture was added (at an MOI of each phage at 10^-1^ to *E. faecalis*), and the cultures incubated. Enrichment culture aliquots were plated on chromogenic agar, and the resultant colony appearance evaluated (Fig. 4). After enrichment of all phage-treated *S. agalactiae–E. faecalis* sets, *S. agalactiae* colonies were dominant on the agar plates. By contrast, in enrichment cultures without phage treatment, only few *S. agalactiae* colonies were observed on the chromogenic agar, while *E. faecalis* colonies were dominant. The same experiment was performed using the modified Lim broth and several *S. agalactiae*–*E. faecalis* combinations (Fig. S4). The data were in agreement with observations made using the GBwSM medium. Thus, the phage mixture treatment improved the *S. agalactiae* culture test by inhibiting the undesirable growth of *E. faecalis*, even when the initial *E. faecalis* cell density was as high 10^5^ CFU/ml.

**FIG 4.**
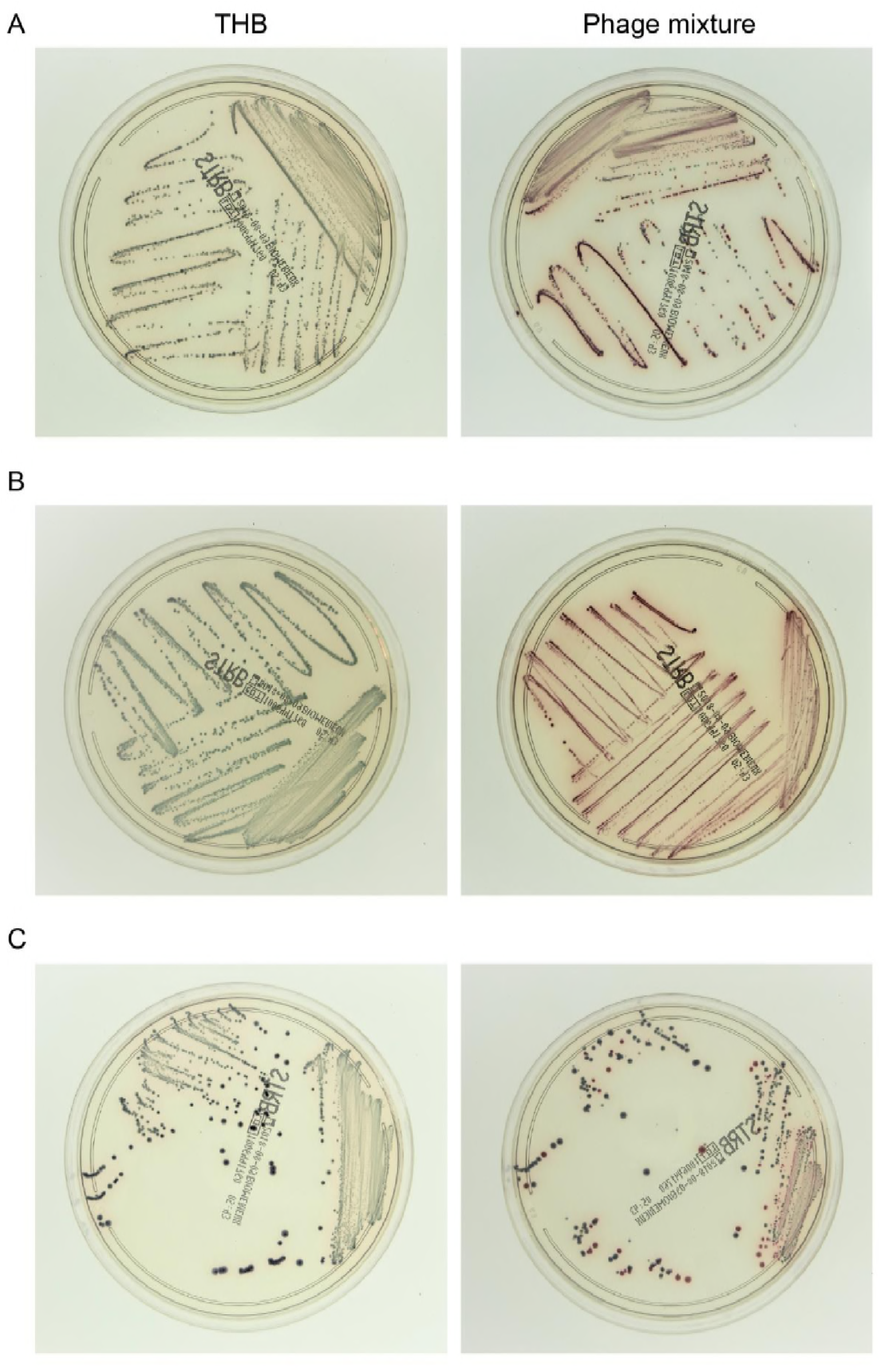
Bacterial identification on chromogenic agar after experimental enrichment co-culture of *S. agalactiae* and *E. faecalis*. Combinations of single strains of *E. faecalis* and *S. agalactiae* were used to inoculate GBwSM, and were cultured in the presence of the phage mixture (MOI of each phage to *E. faecalis:* 10^-1^) or THB. After enrichment culture, aliquots were spread on the chromogenic agar, and the resultant bacterial colonies were evaluated. Colonies of *S. agalactiae* and *E. faecalis* are red and blue, respectively. Left and right panels, photographs of chromogenic agar plates inoculated with enriched cultures treated with THB or phage mixtures, respectively. Representative data for three out of five *S. agalactiae–E. faecalis* sets are shown, namely, KUGBS2–KUEF08 (A), KUGBS1– KUEF24 (B), and KUGBS6–KUEF26 (C).

### Phage application potential in the clinical setting

In the current study, we showed that a specific phage mixture effectively inhibited the growth of *E. faecalis* in an *S. agalactiae* culture test in the experimental setting. Before reagent manufacture and clinical application, phage composition in the phage mixture, the usage per assay (i.e., volume and phage density), storage of the phage mixture, and production cost should be optimized.

As a first consideration, strains insensitive to the phage mixture may occur at a higher than expected rate in the clinical setting. To address this issue, phage sensitivity of *E. faecalis* strain should be constantly examined. The effective spectrum of phages in a mixture can be modified by replacing and/or adding other phages, including newly-isolated phages, naturally-evolved phages, and/or genetically-modified phages (29, 30). E.g., by adding six newly isolated phages to the phage mixture developed in the current study, we increased the inhibition efficiency among the tested *E. faecalis* strains to 96.7% (29/30) (data not shown). Hence, updating the composition of the phage mixture will help to address the problem associated with the phage-insensitive strains.

Next, we considered the usage per assay. Assuming that the volume of the enrichment broth is ca. 3–4 ml, a drop of phage mixture suspension (i.e., 30–50 μl of a solution of ca. 1.0 × 10^7^ PFU/ml of each phage) would suffice for an assay. Such usage per assay is in line with the usage of phage mixture in experiments performed in the current study (Fig. 4). Thus, phage mixture complies with the usage expected in a clinical setting.

Moreover, the quality of the phage mixture should be guaranteed to allow its commercialization. Phages are generally stable at 4°C in culture media for a certain period of time (31, 32), which suggests that phage products can be distributed on a market-scale using the cold chain. Accordingly, we examined the infectious density of each phage solution during storage at 4°C for 270 d, and we did not observe any substantial reduction in phage particle density during this time period (Fig. S5). Shelf life of culture medium is generally up to 6 months (33), and no loss in sensitivity in *S. agalactiae* enrichment broths were observed after at least 4 months (34). Thus, the stability of the phage mixture appeared to be in line with the storage of the enrichment broths.

Finally, the use of phage mixture should be cost-effective. Phage production on a bioreactor-scale has been recently investigated because of increased interest in phage therapy. Consequently, the cost of phage production is estimated to be $4.4 × 10^-13^/phage particle (35-37). Hence, the calculated cost of phage mixture per assay in the current study is only $2.2 × 10^-7^ ($4.4 × 10^-13^/phage particle multiplied by the number of phage particles in the mixture, i.e., 1.0 × 10^7^ PFU/ml × 50 μl). Several commercial phage companies have been founded worldwide (38). Thus, phage mixture may, in theory, be produced for the *S. agalactiae* culture test in a cost-effective manner.

After commercialization of the phage mixture, special attention should be devoted to the biological characteristics of phages to avoid laboratory accidents, since phage products are not commonly used in the clinical microbiology. Phages multiply exponentially and exceed bacterial growth, which might lead to phage contamination and interference with *E. faecalis* culture tests in a clinical laboratory. To avoid such accidents, appropriate safety precautions should be instated when preparing to use phage products. These can include change of gloves after usage, aseptic handling at a specified bench, and storage in a specific cabinet (39). We believe that that the phage mixture satisfies the usage in terms of application volume and density, as long as the knowledge and safety precautions for phage products are disseminated and implemented.

Considering the above, we anticipate that the phage mixture will become commercially available for *S. agalactiae* enrichment broths in the future and will be used in the clinical setting.

## ACKNOWLEDGMENTS

We thank Mr. Takuya Nakajima and Ms. Mio Sasaki (School of Veterinary Medicine, Azabu University, Kanagawa, Japan) for support in performing the experiments. This research was partially supported by research project grants awarded by the Azabu University Research Services Division. All the authors significantly contributed to the work. There are no conflicts of interest to declare.

## REFERENCES

1. Morgan JA, Cooper DB. 2018. Pregnancy, Group B *Streptococcus*. StatPearls Publishing, Treasure Island, FL [updated 11 February, 2018].

2. Edmond KM, Kortsalioudaki C, Scott S, Schrag SJ, Zaidi AK, Cousens S, Heath PT. 2012. Group B streptococcal disease in infants aged younger than 3 months: systematic review and meta-analysis. Lancet 379:547–556.

3. Cagno CK, Pettit JM, Weiss BD. 2012. Prevention of perinatal group B streptococcal disease: updated CDC guideline. Am Fam Physician 86:59–65.

4. Rosa-Fraile M, Spellerberg B. 2017. Reliable detection of group B *Streptococcus* in the clinical laboratory. J Clin Microbiol 55:2590–2598.

5. Dunne WM, Jr., Holland-Staley CA. 1998. Comparison of NNA agar culture and selective broth culture for detection of group B streptococcal colonization in women. J Clin Microbiol 36:2298–2300.

6. Park CJ, Vandel NM, Ruprai DK, Martin EA, Gates KM, Coker D. 2001. Detection of group B streptococcal colonization in pregnant women using direct latex agglutination testing of selective broth. J Clin Microbiol 39:408–409.

7. Binghuai L, Yanli S, Shuchen Z, Fengxia Z, Dong L, Yanchao C. 2014. Use of MALDI-TOF mass spectrometry for rapid identification of group B *Streptococcus* on chromID Strepto B agar. Int J Infect Dis 27:44–48.

8. Baden M, Higashiyama T, Ikemoto T, Okada Y 2016. Evaluation of direct latex agglutination of selective broth for detection of group B streptococcal carriage in pregnant women. J Japn Soc Clin Microbiol 26:7–13.

9. Kurtböke D. 2005. Actinophages as indicators of actinomycete taxa in marine environments. Antonie Van Leeuwenhoek 87:19–28.

10. Muldoon MT, Teaney G, Li J, Onisk DV, Stave JW. 2007. Bacteriophage-based enrichment coupled to immunochromatographic strip-based detection for the determination of *Salmonella* in meat and poultry. J Food Prot 70:2235–2242.

11. Khalifa L, Coppenhagen-Glazer S, Shlezinger M, Kott-Gutkowski M, Adini O, Beyth N, Hazan R. 2015. Complete genome sequence of *Enterococcus* bacteriophage EFLK1. Genome Announc 3:pii:e01308–15.

12. Uchiyama J, Rashel M, Maeda Y, Takemura I, Sugihara S, Akechi K, Muraoka A, Wakiguchi H, Matsuzaki S. 2008. Isolation and characterization of a novel *Enterococcus faecalis* bacteriophage phiEF24C as a therapeutic candidate. FEMS Microbiol Lett 278:200–206.

13. Khalifa L, Gelman D, Shlezinger M, Dessal AL, Coppenhagen-Glazer S, Beyth N, Hazan R. 2018. Defeating antibiotic- and phage-resistant *Enterococcus faecalis* using a phage cocktail *in vitro* and in a clot model. Front Microbiol 9:326.

14. Uchiyama J, Rashel M, Takemura I, Wakiguchi H, Matsuzaki S. 2008. *In silico* and *in vivo* evaluation of bacteriophage phiEF24C, a candidate for treatment of *Enterococcus faecalis* infections. Appl Environ Microbiol 74:4149–4163.

15. Uchiyama J, Takemura I, Satoh M, Kato S, Ujihara T, Akechi K, Matsuzaki S, Daibata M. 2011. Improved adsorption of an *Enterococcus faecalis* bacteriophage phiEF24C with a spontaneous point mutation. PLoS One 6:e26648.

16. de la Rosa M, Perez M, Carazo C, Pareja L, Peis JI, Hernandez F. 1992. New Granada Medium for detection and identification of group B streptococci. J Clin Microbiol 30:1019–1021.

17. Heelan JS, Struminsky J, Lauro P, Sung CJ. 2005. Evaluation of a new selective enrichment broth for detection of group B streptococci in pregnant women. J Clin Microbiol 43:896–897.

18. Nasukawa T, Uchiyama J, Taharaguchi S, Ota S, Ujihara T, Matsuzaki S, Murakami H, Mizukami K, Sakaguchi M. 2017. Virus purification by CsCl density gradient using general centrifugation. Arch Virol 162:3523–3528.

19. Tanizawa Y, Fujisawa T, Nakamura Y. 2018. DFAST: a flexible prokaryotic genome annotation pipeline for faster genome publication. Bioinformatics 34:1037–1039.

20. Tanizawa Y, Fujisawa T, Kaminuma E, Nakamura Y, Arita M. 2016. DFAST and DAGA: web-based integrated genome annotation tools and resources. Biosci Microbiota Food Health 35:173–184.

21. Ruiz-Garbajosa P, Bonten MJ, Robinson DA, Top J, Nallapareddy SR, Torres C, Coque TM, Cantón R, Baquero F, Murray BE, del Campo R, Willems RJ. 2006. Multilocus sequence typing scheme for *Enterococcus faecalis* reveals hospital-adapted genetic complexes in a background of high rates of recombination. J Clin Microbiol 44:2220–2228.

22. Jolley KA, Maiden MC. 2010. BIGSdb: scalable analysis of bacterial genome variation at the population level. BMC Bioinformatics 11:595.

23. Kumar S, Stecher G, Tamura K. 2016. MEGA7: molecular evolutionary genetics analysis version 7.0 for bigger datasets. Mol Biol Evol 33:1870–1874.

24. Vijayakumar PP, Muriana PM. 2015. A microplate growth inhibition assay for screening bacteriocins against *Listeria monocytogenes* to differentiate their mode-of-action. Biomolecules 5:1178–1194.

25. Kanda Y 2013. Investigation of the freely available easy-to-use software “EZR” for medical statistics. Bone Marrow Transplant 48:452–458.

26. Lefkowitz EJ, Dempsey DM, Hendrickson RC, Orton RJ, Siddell SG, Smith DB. 2018. Virus taxonomy: the database of the International Committee on Taxonomy of Viruses (ICTV). Nucleic Acids Res 46:D708–D717.

27. Klumpp J, Lavigne R, Loessner MJ, Ackermann HW. 2010. The SPO1-related bacteriophages. Arch Virol 155:1547–1561.

28. Nes IF, Diep DB, Holo H. 2007. Bacteriocin diversity in *Streptococcus* and *Enterococcus.* J Bacteriol 189:1189–1198.

29. Ando H, Lemire S, Pires DP, Lu TK. 2015. Engineering modular viral scaffolds for targeted bacterial population editing. Cell Syst 1:187–196.

30. Koskella B, Brockhurst MA. 2014. Bacteria-phage coevolution as a driver of ecological and evolutionary processes in microbial communities. FEMS Microbiol Rev 38:916–931.

31. Ackermann H, Tremblay D, Moineau S. 2004. Long-term bacteriophage preservation WFCC Newsl 38:35–40.

32. Lobocka MB, Glowacka A, Golec P. 2018. Methods for bacteriophage preservation. Methods Mol Biol 1693:219–230.

33. Ulisse S, Peccio A, Orsini G, Di Emidio B. 2006. A study of the shelf-life of critical culture media. Vet Ital 42:237–247.

34. Carvalho Mda G, Facklam R, Jackson D, Beall B, McGee L. 2009. Evaluation of three commercial broth media for pigment detection and identification of a group B *Streptococcus (Streptococcus agalactiae).* J Clin Microbiol 47:4161–4163.

35. Agboluaje M, Sauvageau D. 2018. Bacteriophage production in bioreactors. Methods Mol Biol 1693:173–193.

36. Krysiak-Baltyn K, Martin GJO, Gras SL. 2018. Computational modeling of bacteriophage production for process optimization. Methods Mol Biol 1693:195218.

37. Krysiak-Baltyn K, Martin GJO, Gras SL. 2018. Computational modelling of large scale phage production using a two-stage batch process. Pharmaceuticals (Basel) 11:pii:E31.

38. Forde A, Hill C. 2018. Phages of life - the path to pharma. Br J Pharmacol 175:412-418.

39. Los M, Czyz A, Sell E, Wegrzyn A, Neubauer P, Wegrzyn G. 2004. Bacteriophage contamination: is there a simple method to reduce its deleterious effects in laboratory cultures and biotechnological factories? J Appl Genet 45:111–120.

